# MDSC depletion during immunization with heat-killed *Mycobacterium tuberculosis* increases protection against BCG infection

**DOI:** 10.1101/2025.01.31.635639

**Authors:** Arpa Aintablian, Haisam Alattar, Laura Cyran, Christoph Schoen, Nelita Du Plessis, Gerhard Walzl, Ulrich Schaible, Natalie E. Nieuwenhuizen, Manfred B. Lutz

## Abstract

Tuberculosis (TB) is still one of the deadliest infectious diseases globally. Although i.d. BCG immunization offers limited protection, a vaccine based on *Mycobacterium tuberculosis* (Mtb) has yet to be approved. Our previous findings demonstrated that s.c. immunization with heat-killed Mtb significantly increased the number of monocytic myeloid-derived suppressor cells (M-MDSC) in mice. Therefore, we hypothesized that the defense against a subsequent BCG infection would be impaired in Mtb-immunized mice. Surprisingly, mice vaccinated with Mtb were protected against a BCG infection and showed elevated frequencies and activation of DC and mycobacteria-specific T cells despite high frequencies and suppressor activity of M-MDSC. Genetic ablation of CCR2^+^ monocytic cells or pharmacological intervention with all-trans retinoic acid (ATRA) reduced the frequency of Mtb-induced M-MDSC, enhanced frequencies, and activation of dendritic cells (DC) and CD4^+^ T cells, and resulted in decreased bacterial loads in the lung and spleen. These findings offer fresh perspectives on TB vaccination using heat-killed Mtb despite parallel unwanted vaccine-induced M-MDSC. M-MDSC depletion by ATRA further tips the balance towards immunity and should be considered an adjunct host-directed therapy with TB vaccines in humans.

## Introduction

TB, an infectious disease caused by Mtb, still accounts for a significant burden of annual morbidity and mortality worldwide. Mtb-based vaccine candidates have been shown to induce T cell responses but have so far failed to lead to an approved vaccine ^1^. Currently, multiple Mtb-based vaccine candidates are in various phases of clinical trials ^2^. Bacille-Calmette-Guérin (BCG), the live attenuated form of *M. bovis* (Mbov), which causes TB in cattle, is the only available FDA-approved TB vaccine to date. In many countries, BCG is routinely administered to infants shortly after birth, effectively preventing meningeal and miliary TB and displaying 60-80% protective efficacy against pulmonary disease in children ^3^. However, the BCG vaccine shows limited and highly variable protective efficacy in adults, representing the most active TB cases ^2,4^. While vaccines are typically generated from the same microbe, the BCG vaccine represents a heterologous vaccine, lacking numerous antigenic determinants that are present in Mtb ^5^. However, since Mtb has developed many immune evasion mechanisms preventing immune responses, especially immune memory, BCG appears to represent the best compromise for a TB vaccine so far ^6^. The challenge remaining is to apply Mtb as a vaccine but circumvent critical immune evasion mechanisms preventing the immune memory and, thereby, the vaccine success.

As a novel immune evasion mechanism of Mtb, we found an accumulation of myeloid-derived suppressor cells (MDSC) in the blood of TB patients and healthy individuals recently exposed to Mtb ^7^. Similarly, suppressive MDSC were detected in the lungs of Mtb-infected mice, providing a suitable environment for pathogen survival ^8^. Two major MDSC subsets have been classified as CD11b^+^ Ly6C^high^ Ly6G^-^ M-MDSC and CD11b^+^ Ly6C^low^ Ly6G^+^ granulocytic MDSC (G-MDSC, also known as polymorphonuclear (PMN)-MDSC), determined by their monocytic or granulocytic hematopoietic origin, respectively ^9–12^. MDSC possess different immunosuppressive mechanisms. These include T cell receptor (TCR) nitrosylation, attributed to their expression of intracellular inducible nitric oxide synthase (iNOS) and consequent release of nitric oxide (NO), and T cell metabolic starvation due to arginase 1 (Arg1)-dependent catabolism of arginine from the extracellular space ^13,14^.

Our previous work showed that even a heat-killed Mtb double vaccine massively induced M-MDSC ^15^. These M-MDSC accumulated in the spleen and displayed potent DC-killing capacity in an iNOS-dependent manner *in vivo*, thereby indirectly suppressing effector T cell responses ^15^. It remains unknown whether a heat-killed BCG or *Mycobacterium bovis* (Mbov) vaccine also induce the differentiation of monocytes into suppressive M-MDSC, resulting in insufficient protection. In fact, few reports exist about the protective innate and/or adaptive immune effects of BCG vaccination on a subsequent Mbov infection in mice ^16^. Despite differences among mycobacterial species, most immune effects against challenges have been described with heterologous Mtb for mice, while BCG vaccination has been tested frequently against Mbov infection in cows. However, in cattle, immune responses were scarcely followed due to the limited availability of reagents. Most experimental studies showed a reduced disease severity, but field trials also showed protection from infection ^17,18^.

Host-directed therapies targeting MDSC against tumors show promising results in animal models and clinical trials with cancer patients ^19,20^. Since chronic infectious diseases such as HIV or TB still suffer from the lack of effective vaccines, the extrapolation of host-directing therapies targeting MDSC are also discussed for these infections ^21^. All-trans retinoic acid (ATRA), an active metabolite of vitamin A, has been shown to interfere with MDSC development by shifting it towards pro-inflammatory myeloid effector cells ^22^. Promising outcomes have been observed in murine Mtb infection models ^8,23–25^, and also in cancer patients ^26,27^. Since ATRA was also successfully combined with cancer vaccines in patients ^28^ but has not yet been tested in combination with an Mtb vaccine, we applied it here in our mouse model.

To gain further insights into the conditions favoring vaccine-induced M-MDSC by mycobacteria, we altered multiple factors in an established immunization protocol ^15^, including the prime-boost immunization interval, type of adjuvant, vaccine dose, and method of Mtb preparation, as well as the use of different dead bacterial species within the vaccine. We also investigated the effect of heat-killed Mtb and BCG vaccines on subsequent live BCG infection in mice. Our results indicate that Mtb appears quantitatively superior to other bacteria tested here in the induction of M-MDSC. Surprisingly, an intranasal challenge of Mtb-immunized mice with BCG infection resulted in simultaneous iNOS-dependent M-MDSC-mediated suppression and effector immune responses by DC and T cells in the lung, biased towards protective effects. Genetic CCR2-targeted depletion or pharmacological interference by ATRA with M-MDSC reduced their frequencies, enhanced DC and CD4^+^ T cell frequencies, and further reduced bacterial burdens in the lung and spleen. These results provide new insights into the relationship between M-MDSC-driven suppression and DC/T cell-mediated immunity. Combining Mtb vaccination with ATRA treatment may be considered a new strategy for developing an effective Mtb-based TB vaccine in humans.

## Results

### Mtb outcompetes Msm and Listeria in M-MDSC induction, in contrast to BCG, which fails to induce M-MDSC

Our previous findings indicated that double immunization with heat-killed Mtb in IFA resulted in potent systemic M-MDSC induction in mouse spleens ^15^. Here, we examined M-MDSC induction by double immunization with different bacteria in IFA, including heat-killed BCG, Msm, List, wild-type Mbov, and single vaccination with Mtb followed by List (Figure 1A). As observed before, splenomegaly occurred after 2xMtb immunization, which was also seen here after immunization with Mtb/List, but not with the other double vaccines (Figure 1B).

**Figure 1:**
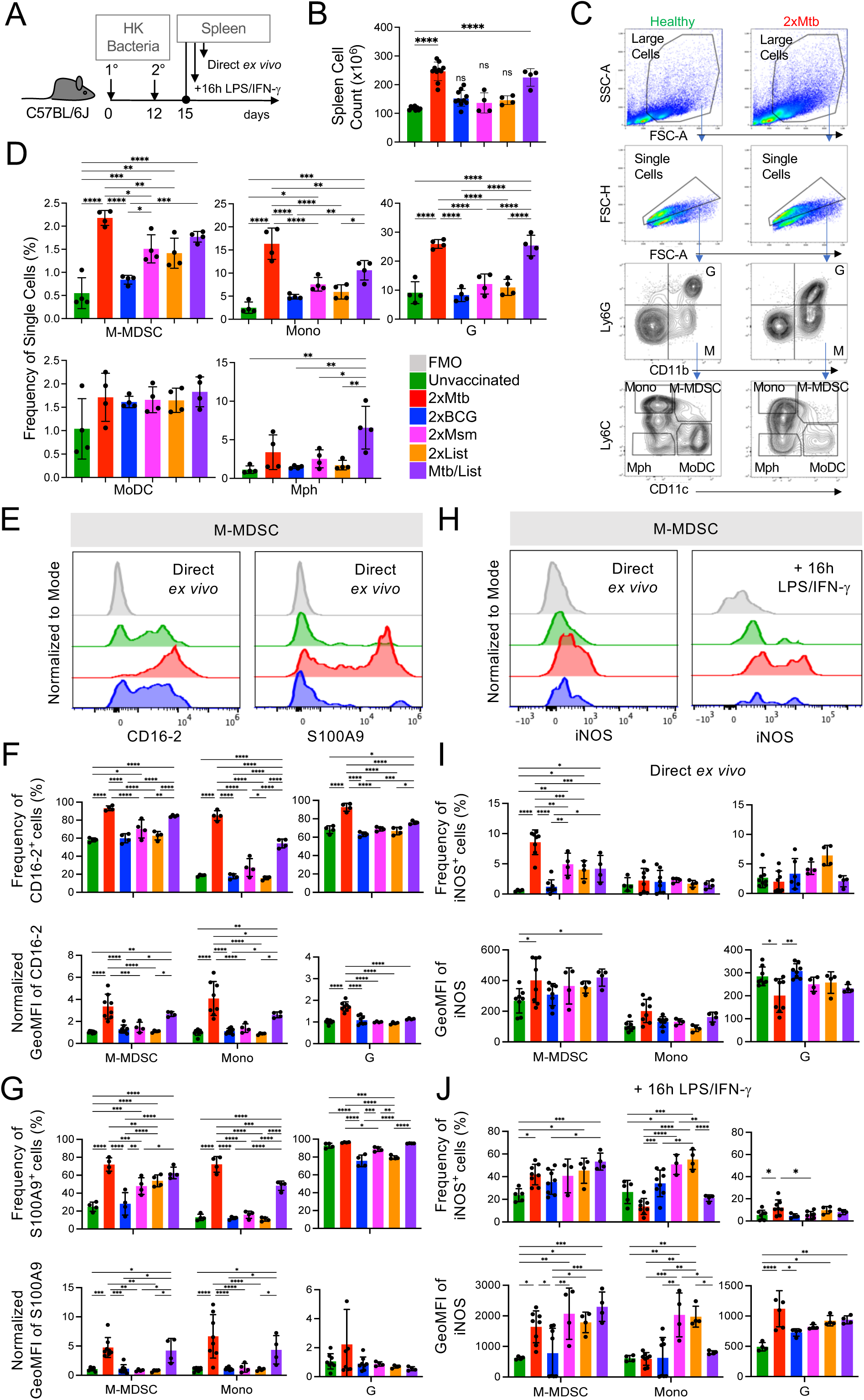
Superior potential of 2xMtb immunization over other bacteria to increase M-MDSC frequencies, activation and iNOS expression. **A.** Time scale of the standard protocol for primary and secondary immunization. Mice remained unvaccinated or were immunized s.c. with 10^6^ CFU of the indicated heat-killed (HK) bacteria in IFA at day 0, followed by a booster dose at day 12. Spleens were harvested and analyzed at day 15. **B.** Spleen cell numbers obtained from the indicated immunization conditions. **C.** Gating strategy to identify granulocytes (G), M-MDSC, monocytes (Mono), macrophages (Mph), and monocyte-derived DC (MoDC). **D.** Frequencies of splenic cell types indicated in C. **E.** Representative histograms showing expression of CD16-2 and S100A9 activation markers in M-MDSC gate. **F, G.** Frequency and normalized GeoMFI of CD16-2 and S100A9 in M-MDSC, Mono and G gates, respectively, directly *ex vivo***. H.** Spleen cells were restimulated *in vitro* by LPS/IFN-γ for 16h then stained for iNOS. Histograms showing iNOS expression directly *ex vivo* and after restimulation *in vitro* in M-MDSC gate. **I, J.** Frequency and GeoMFI of iNOS in M-MDSC, Mono and G gates directly *ex vivo* (I) and after restimulation (J). **B, D:** Statistics by ordinary one-way ANOVA with multiple comparisons and Tukey’s post test. n=4-10 biological replicates, 2-4 independent experiments. **F, G, I, J:** Statistics by two-way ANOVA with multiple comparisons and Tukey’s post test. n=3-8 biological replicates, 2-4 independent experiments. * p<0.05; ** p<0.01; *** p<0.001; **** p<0.0001

Spleens were further analyzed by flow cytometry for monocytic and granulocytic cells, which could be distinguished according to their relative surface expression of CD11b and Ly6G. Monocytic cells were further subdivided into Ly6C^high^ CD11c^-^ classical monocytes (Mono), Ly6C^high^ CD11c^+^ M-MDSC, Ly6C^-^ CD11c^+^ monocyte-derived DC (MoDC), and Ly6C^-^ CD11c^-^ macrophages (Mph) (Figure 1C), as described earlier. Frequencies of M-MDSC, Mono, and granulocytic cells (G), almost exclusively neutrophils, dramatically increased with the 2xMtb and Mtb/List vaccines, while only to a lower extent with 2xMsm and 2xList (Figure 1D). Interestingly, 2xBCG immunization did not significantly increase the spleen cellularity and frequencies of M-MDSC, classical monocytes, and granulocytic cell (G) frequencies compared to untreated controls (Figure 1B, D). Mph and MoDC showed only marginal differences between the groups (Figure 1D) and were not followed further.

To further characterize the activation state of the myeloid cell subsets, we used CD16-2 and S100A9 as M-MDSC activation markers. CD16-2 is the activating FcγRIV ^29^, and S100A9 belongs to the family of alarmins ^30^, which are both constitutively expressed on neutrophils and G-MDSC but not monocytes. 2xMtb and Mtb/List immunizations strongly induced the expression of CD16-2 and S100A9 by M-MDSC, Mono, and G populations, compared to vaccines from all other bacterial species (Figure 1E-G). Conversion of monocytes into suppressive MDSC with NO-releasing capacity is a two-step process that first requires an initial ‘licensing’ and expansion step, e.g., by GM-CSF or bacterial immunization. Upon secondary challenge, e.g., by LPS/IFN-γ, functional suppression is activated by upregulation of iNOS and release of NO. Therefore, we analyzed iNOS expression directly and after re-stimulation of splenocytes *in vitro* with LPS/IFN-γ. Immunization with 2xMtb resulted in a strong increase in the frequency of iNOS^+^ M-MDSC, whereas 2xBCG did not, and 2xMsm, 2xList, and Mtb/List moderately increased the frequencies (Figure 1H, I). Re-stimulation with LPS/IFN-γ strongly increased iNOS expression levels and frequencies of iNOS^+^ M-MDSCs in most immunization groups. Still, the expression levels of iNOS remained low in cells from BCG-immunized mice (Figure 1H, J).

Due to the unique ability of 2xMtb to induce M-MDSC, we tested how the bacterial dose, the immunization interval, or the adjuvant type influenced the myeloid cell compartment and M-MDSC induction. Immunization of mice with 100-fold lower Mtb doses reduced spleen cell numbers and frequencies of classical monocytes but did not affect the frequency of M-MDSC and granulocytic cells compared to our standard immunization protocol (Supplementary Figure 1A, D). Application of the booster vaccine after longer intervals (Supplementary Figure 1B, E) or the use of alum instead of IFA (Supplementary Figure 1C, F) had no significant effect on spleen cell numbers or M-MDSC and granulocyte frequencies but resulted in lower numbers of monocytes compared to our standard immunization protocol. To investigate whether the method of Mtb preparation or the attenuation of BCG affects M-MDSC induction, we compared vaccines prepared from heat-killed and lyophilized Mtb to those prepared with heat-killed but non-lyophilized Mtb (MtbM). Also, comparisons were made between BCG and its non-attenuated counterpart, Mbov. Vaccination with 2xMtbM led to lower frequencies of monocytes and granulocytic cells than 2xMtb but similar or only slight reductions in spleen cell counts and M-MDSC frequencies and activation markers (Supplementary Figure 2A-H). 2xMbov immunization resulted in similar frequencies of M-MDSCs to 2xBCG and was comparable in most readouts apart from higher levels of CD16-2 on granulocytic cells and iNOS+ on directly analyzed M-MDSCs (Supplementary Figure 2A-H). This indicates that neither the Mtb preparation nor the attenuation of wild-type Mbov to BCG has a major influence on the capacity of these strains to induce M-MDSC. Together, these results suggest that intrinsic components of the Mtb bacterial strain may be responsible for inducing M-MDSC.

### Mtb immunization before BCG infection induces lung M-MDSCs but reduces lung bacterial loads, in contrast to BCG vaccination

Next, we sought to investigate the influence of 2xMtb or 2xBCG immunizations, which contrasted in their capacity to induce M-MDSCs, on a subsequent BCG infection. Mice were immunized accordingly or remained untreated and were infected i.n. with live BCG three weeks after the booster vaccination (Figure 2A). There was a reduction in bacterial load in the lung but an increase in the spleen after 2xMtb immunization compared to unvaccinated mice (Figure 2B). Surprisingly, 2xBCG vaccinated animals showed a similar increase in bacterial load in the spleen and a non-significant trend for higher bacterial load in the lung (Figure 2B). As observed in 2xMtb immunized animals ^15^, visible splenomegaly also remained after infection with BCG, correlating with increased cell counts in the spleen and lung (Figure 2C). The unexpected protective effect of the 2xMtb immunization despite M-MDSC induction was confirmed by the anatomical examination of the lungs, with 2xMtb vaccinated animals showing clearer lungs with fewer patches of inflammation (Figure 2D). 2xMtb immunization before BCG infection enhanced the frequencies of M-MDSC, monocytic, and granulocytic cells in the lung (Figure 2E). Myelopoiesis induced by 2xMtb immunization persisted after BCG infection, shown by increased numbers of M-MDSC, monocytes, and granulocytic cells in the spleen (Figure 2E).

**Figure 2:**
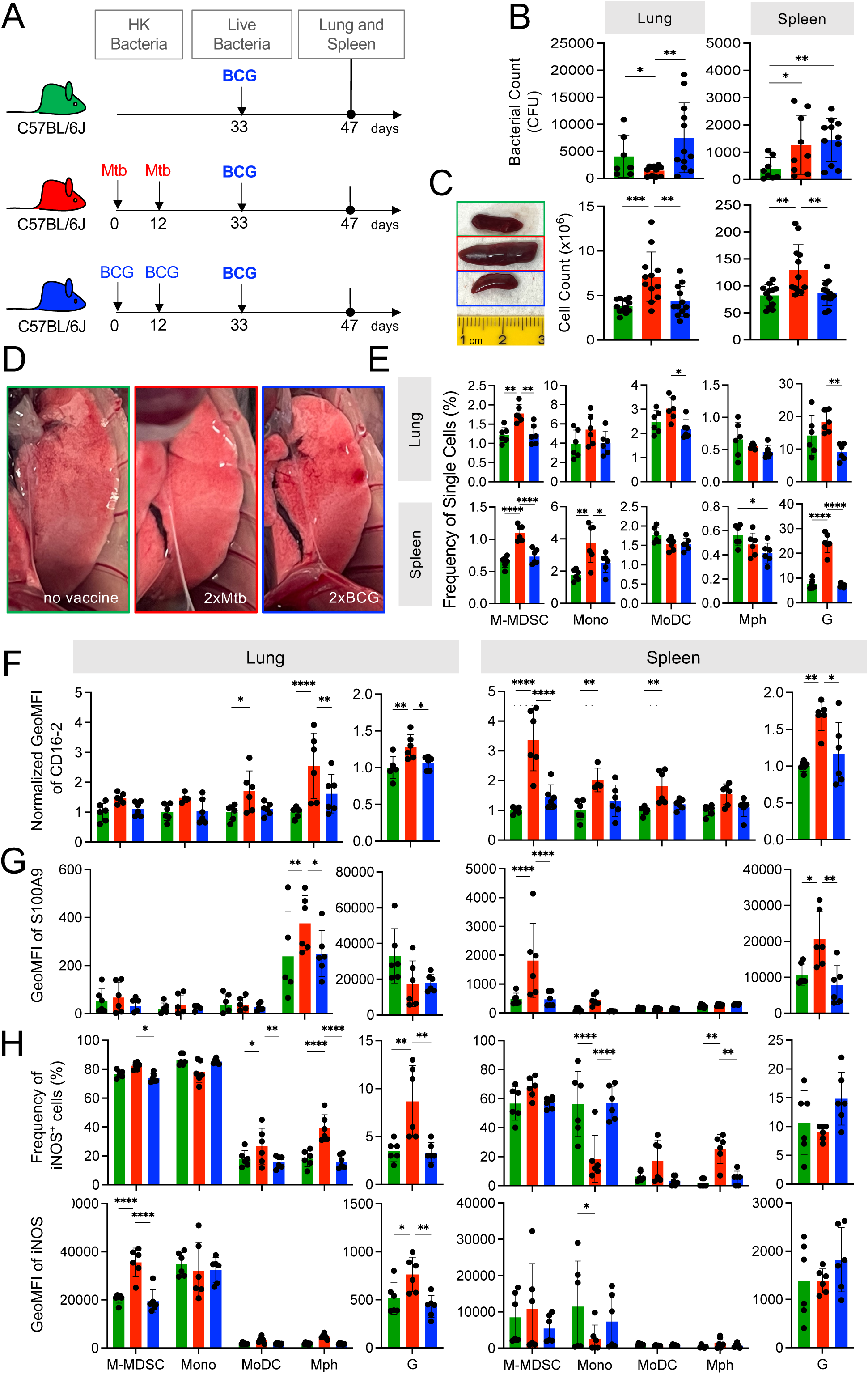
BCG-infected lungs of Mtb- but not BCG-immunized mice show increased lung infiltration of M-MDSC, but they lose activation and suppression markers. **A.** Scheme showing immunization protocols with subsequent live BCG infection. Mice remained untreated or were immunized s.c. with 10^6^ CFU heat-killed (HK) bacteria in IFA at day 0 followed by a booster dose at day 12. On day 33, all mice were infected i.n. with 10^7^ CFU live BCG; lungs and spleens were harvested and analyzed at day 47. **B.** Bacterial loads in whole lung and spleen. **C.** Representative spleen photographs and cell numbers of lung and spleen as indicated. **D.** Representative photographs of 12 lungs per group showing showing reduction in areas of inflammation (red/white patches) in 2xMtb-immunized mice. **E.** Frequencies of myeloid cells in specified gates of lung and spleen. Gates as in Fig. 1C. **F, G.** Expression of CD16.2 and S100A9 by the indicated cell types via flow cytometry. **H.** Lung and spleen cells were restimulated *in vitro* by LPS/IFN-γ for 16h then stained for iNOS. **B:** Statistics by unpaired, two-tailed t-test. n=8-10 biological replicates, 2 independent experiments. **C, E-H:** Statistics by ordinary one-way ANOVA with multiple comparisons and Tukey’s post test. Data from n=6 mice with 2 technical replicates from 2 independent experiments. * p<0.05; ** p<0.01; *** p<0.001; **** p<0.0001

Flow cytometric analysis of the spleen after BCG infection revealed two cell populations with differential Ly6G and CD11b expression levels within the G gate, which we designated G^high^ and G^low^ (Supplementary Figure 3A). Cells in the G^high^ gate expressed higher levels of S100A9 and CD16-2 activation markers than those in the G^low^ gate, and only moderate differences appeared in the frequencies and expression levels of iNOS and Arg1 suppression markers (Supplementary Figure 3B, C). Overall, the cells in the G^high^ and G^low^ gates of the Mtb-immunized group appear to reflect differentiation or activation states, with the G^high^ cells being more functionally advanced than the G^low^ population. These data agree with our previous findings, where the G^high^ and G^low^ cells reflected mature and immature neutrophils in the spleen, respectively, as judged by their nuclear shape ^31^. Thus, the total G gate was used for further analyses. We also investigated the activation and functional state of alveolar macrophages (AM) in our model, as they are the first immune cells to encounter mycobacteria in the lung and are critical in determining disease outcome ^32^. The CD11c^+^ fraction within the Ly6G^-^ CD11b^-^ (dn) gate was designated as AM (Supplementary Figure 3D, E). The CD11b^-^Ly6G^-^ G^low^ population in the lung expressed low levels of CD11c and showed a dispersed forward and side scatter profile. Hence, they did not appear to represent immature neutrophils as in the spleen (Supplementary Figure 3F). Thus, only the G^high^ populations were analyzed in the lungs. Analyses of AM’s frequency, activation, and suppression markers showed no differences between immunization groups (Supplementary Figure 3G, H, I). This agrees with recent findings that mycobacterial infection switches from AM to monocyte-derived Mph after a few days ^33^.

We then studied the activation state of the monocytic and granulocytic cells induced by live BCG infection directly *ex vivo*. While 2xMtb immunizations before BCG infection resulted in up-regulation of the activation markers CD16-2 and S100A9 and down-regulation of iNOS in splenic M-MDSC and monocytes, this was not observed in the lung (Figure 2F-H). Instead, Mph, MoDC, and granulocytes showed signs of activation (Figure 2F-G) and increased frequencies of iNOS^+,^ especially in the lung after *in vitro* restimulation (Figure 2H). Mph and MoDC showed no improvement of iNOS expression as indicated by GeoMFI in contrast to M-MDSC and granulocytes (Figure 2H). The activation and expansion of iNOS^+^ Mph and granulocytes may indicate that they achieved immunogenic properties since it has been shown that intracellular killing of mycobacteria by macrophages is iNOS dependent ^34^. This may point to a switch from a dominant suppressive myeloid compartment towards immunogenic but iNOS^-^ myeloid cell types.

Together, 2xBCG immunizations did not protect against BCG infection and remained without induction of immunity; rather, they tended to enhance lung bacterial growth and did not affect the frequencies of monocytic and granulocytic cells. In contrast, an equivalent amount of 2xMtb immunizations led to lower bacterial burdens and inflammation in the lung after BCG infection and induced both M-MDSC and myeloid effector cells. These results suggest that 2xMtb vaccination may induce robust immunity against BCG despite the parallel M-MDSC induction.

### Expansion and activation of DC subsets in the lungs of Mtb-immunized mice

To corroborate the hypothesis that live BCG infection after 2xMtb vaccination triggers a switch towards immunogenic myeloid cell phenotypes, we investigated the responses of myeloid cells, including DC subsets, after 2xMtb or 2xBCG immunizations and live BCG infection. The gating strategy was adopted from our previous work ^35^ and allowed us to separately analyze pDC, MoDC, cDC1, cDC2, and XCR1/CD11b-double negative (DN) DC, and different Mph subsets (CD11c^high^ CD11b^-^ in the spleen and CD11c^high^ CD11b^-^ AM in the lung) (Figure 3A). 2xMtb immunizations followed by BCG infection resulted in enhanced frequencies of cDC1, MoDC, and DN populations in the spleen and lung, and also cDC2 in the spleen, while 2xBCG immunization followed by BCG infection showed no effect (Figure 3B).

**Figure 3:**
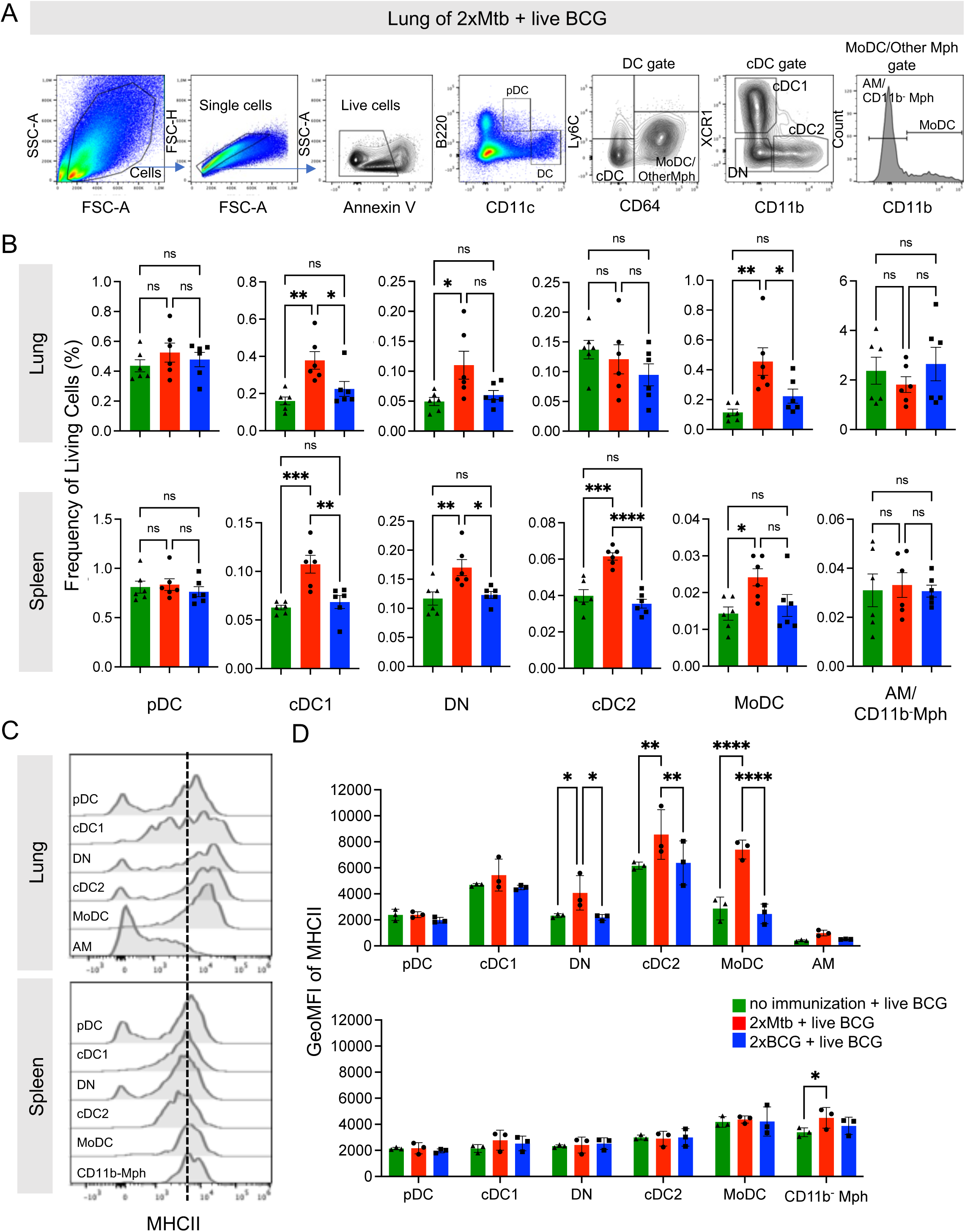
BCG-infected lungs of 2xMtb-immunized mice show high activation of several dendritic cell and monocyte subsets. **A.** Gating strategy to identify DC subsets. Plasmacytoid DC (pDC) were defined as as B220^+^ CD11c^low^. CD11c^hi^ DC were divided into Ly6C^-^ CD64^+^ Monocyte-derived DC (MoDC), other Mph and Ly6C^-^ CD64^-^ conventional DC (cDC). The Monocyte-derived DC (MoDC) and other Mph gate was further subdivided into CD11b^+^ MoDC, and CD11b^-^ AM (in lung) or CD11b^-^ Mph (in spleen). The cDC gate was further subdivided into XCR1^+^ CD11b^-^ cDC1 and XCR1^-^ CD11b^+^ cDC2. Cells negative for XCR1 and CD11b were gated as double negative DC (DN). **B.** Frequencies of living cells in specified gates of lung and spleen. Gates as in (A)**. C.** Histogram overlays showing MHCII expression of indicated cell subsets in lung and spleen. **D.** GeoMFI of MHC II in indicated gates of lung and spleen. Gates as in A. **B:** Statistics by ordinary one-way ANOVA with multiple comparisons and Tukey’s post test. n=12 biological replicates pooled 2 by 2 (therefore n=6), 2 independent experiments. **D:** Statistics by ordinary two-way ANOVA with multiple comparisons and Tukey’s post test. n=6 biological replicates pooled 2 by 2 (therefore n=3). Not significant (ns); * p<0.05; ** p<0.01; **** p<0.0001. Dotted lines separate MHC II^low^ from MHC II^high^ expression for immature and mature DC subsets, respectively.

MHC II expression of the DC was further used as an activation marker (Figure 3C, D). In the lung, 2xMtb immunizations induced the highest MHC II expression in MoDC, cDC2, and DN populations (Figure 3C, D) and moderate levels in cDC1. In contrast, no MHC II expression was observed in the spleen, except moderately in CD11b^-^ Mph under 2xMtb immunization condition (Figure 3C, D). Again, no DC activation was observed when mice were immunized with 2xBCG and infected with live BCG in the lung and spleen (Figure 3C, D). These data point towards the activation of DC subsets in BCG-infected animals that received prior 2xMtb but not 2xBCG vaccination, suggesting a greater ability to activate protective T cell responses.

### Mtb immunization before BCG infection results in enhanced lung CD4^+^ and CD8^+^ T cell responses

It remained to be determined how the mixed induction of immunostimulatory and suppressive myeloid cells would translate into adaptive immunity. To test this, antigen-specific versus polyclonal endogenous T cell responses were investigated in the 2xMtb- or 2xBCG-vaccinated and BCG-infected mice. Mice received CD45.1^+^ splenocytes from P25 mice carrying a TCR-transgene in CD4^+^ T cells with specificity for peptide 25 of mycobacteria Antigen 85B ^36^. This enabled us to follow the peptide-specific T cell expansion against Mtb and BCG and compare it to the endogenous T cell repertoire (Figure 4A). A massive expansion of antigen-specific P25 and endogenous CD4^+^ T cells was observed in the lungs of Mtb-immunized but not BCG-immunized mice infected with BCG (Figure 4B). CD8^+^ T cell populations showed no change in immunized mice compared to unvaccinated controls. To test the P25 and endogenous T cell proliferation, bulk lung, and spleen cells were re-stimulated *in vitro* with peptide P25 for 72h, followed by assessment of the Ki67^+^ proliferating cells. P25 T cells proliferated heavily only in the 2xMtb vaccinated animals. In contrast, T cell responses of animals that received 2xBCG were not elevated and remained at a very low level, similar to those of unvaccinated mice (Figure 4C). Significantly increased proliferation of CD8^+^ and CD4^+^ endogenous T cells could also be detected in the lung, but not spleen, in mice after 2xMtb immunization (Figure 4C). Interestingly, P25 T cell proliferation in the lung was further enhanced by adding an iNOS inhibitor (L-NMMA) but not an Arg1 inhibitor (nor-NOHA) to the culture, suggesting the presence of iNOS-mediated suppressive activity. 2xBCG immunization followed by BCG infection only increased CD8^+^ T cell proliferation upon such *in vitro* re-stimulation (Figure 4C).

**Figure 4:**
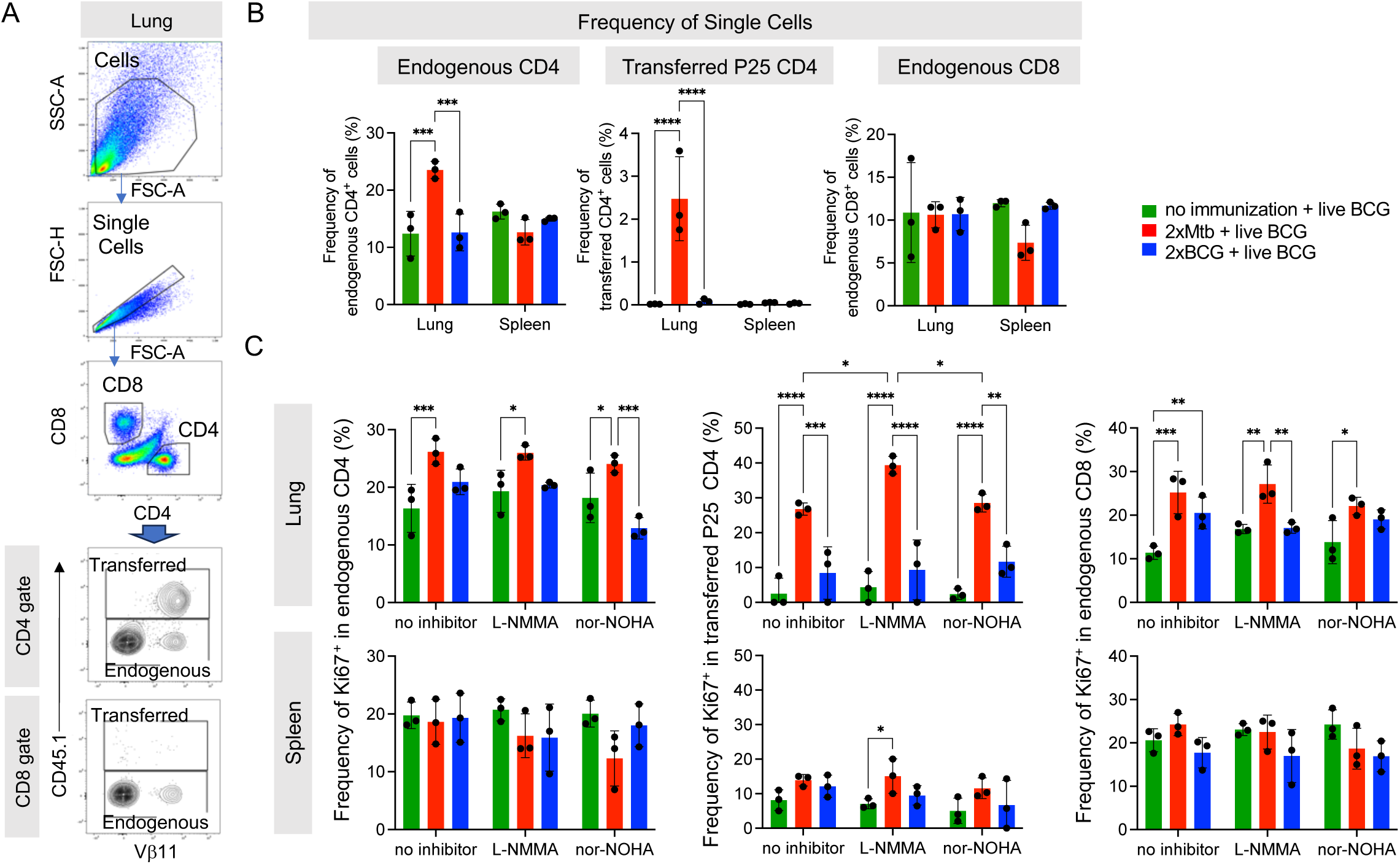
Enhanced CD4^+^ and CD8^+^ T cell immune response in the lungs of 2xMtb-immunized and BCG-infected mice. **A.** Representative lung flow cytometry plots and gating strategy to identify endogenous and transferred CD4 and CD8 T cells. Initial gating included FSC-A/SSC-A and doublet exclusion. CD4^+^ and CD8^+^ lymphocytes were segregated based on their surface expression of CD4 or CD8, respectively. Transferred CD4^+^ transgenic P25 T cells were gated as CD45.1^+^ based on their expression of the congenic marker. Endogenous CD4^+^ and CD8^+^ T cells were gated as negative for CD45.1. **B.** Frequencies of single cells in lung and spleen endogenous CD4, transferred P25 CD4 and endogenous CD8 gates. Gates as in A. **C.** Bulk lung and spleen cells were stimulated at day 47 with P25 peptide and cultured for 3 days with or without iNOS inhibitor L-NMMA or Arg1 inhibitor nor-NOHA. T cell proliferation was measured by flow cytometry and is depicted as frequency of Ki67^+^ within endogenous CD4, transferred CD4 and endogenous CD8 T cell subsets. Statistics with two-way ANOVA with multiple comparisons and Tukey’s post test. n=6 biological replicates pooled 2 by 2 (therefore n=3). * p<0.05, ** p<0.01, *** p<0.001, **** p<0.0001.

These findings indicate mycobacteria-specific P25 and endogenous CD4^+^ T cell expansion in the lung as the major infected organ after 2xMtb immunizations, followed by BCG infection. The further enhanced proliferation of T cells from the lung after adding the iNOS inhibitor follows the decreased M-MDSC frequency and activation observed in these mice.

### Depletion or inhibition of M-MDSC before BCG infection further lowers bacterial loads in the lung and enhances cDC1 frequencies and T cell responses

Re-stimulation of mycobacteria-specific P25 T cells from the lung indicated that M-MDSC were still partially suppressing the T cell response *in vitro*. To demonstrate that M-MDSC are also active *in vivo,* we used *Ccr2*-DTR-CFP mice, where all CCR2^+^ monocytes and monocyte-derived cells can be depleted by injection of DT. We vaccinated with 2xMtb and performed the depletion after M-MDSC induction but before the BCG infection or after the BCG infection (Figure 5A). Depleting CCR2^+^ cells after M-MDSC induction (days 15, 16, 17) further reduced the bacterial loads in the lung and spleen, although this did not reach statistical significance in the spleen (Figure 5B). In contrast, depletion of CCR2^+^ cells after BCG infection at the peak of cDC activation and subsequent T cell response (days 34,35,36) did not further reduce the bacterial loads in the lung or spleen, similar to DT injections as a control into WT mice (Figure 5B).

**Figure 5:**
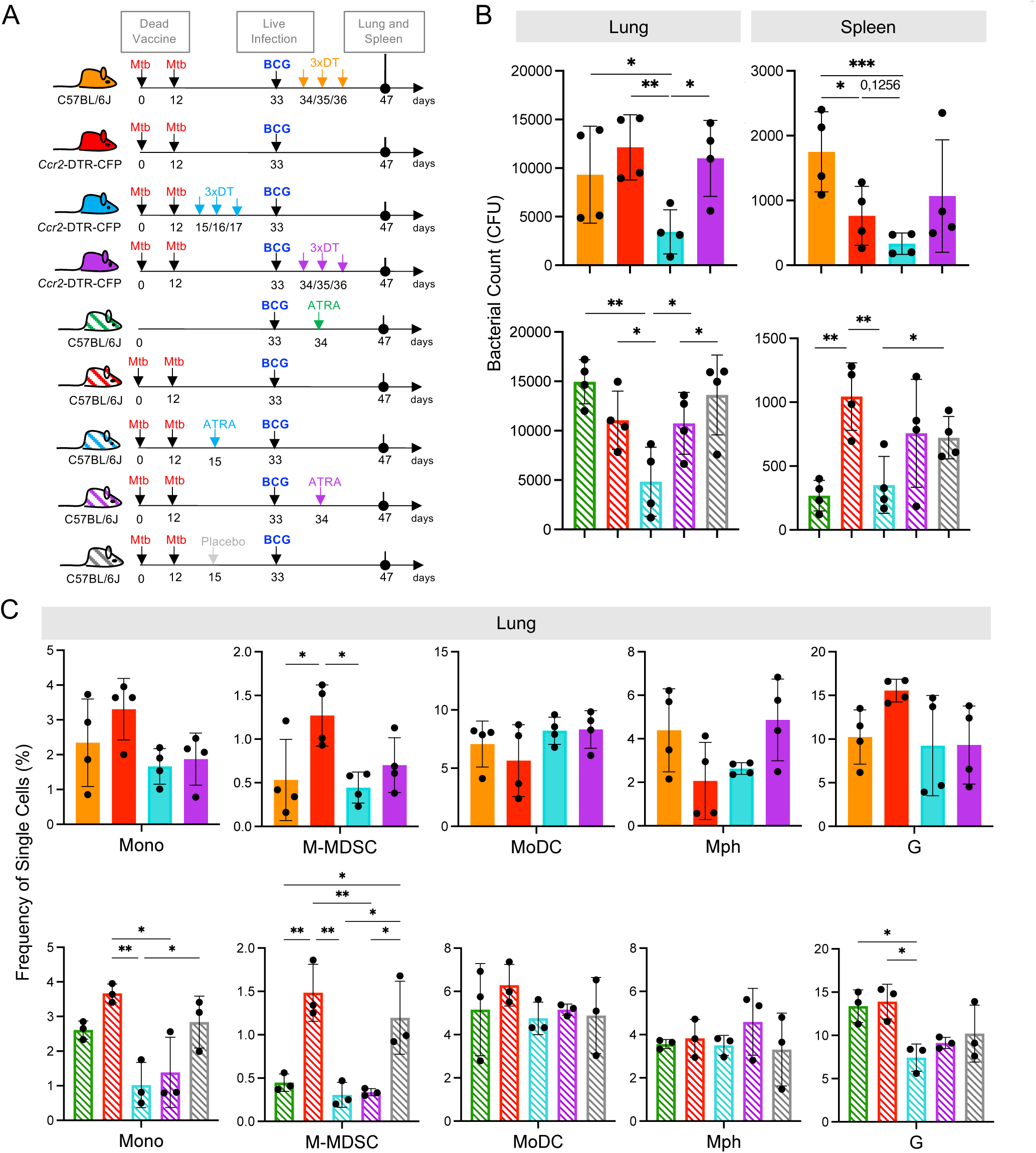
Diphteria toxin (DT) or All-trans retinoic acid (ATRA) treatment prior to intranasal BCG-infection further decreases lung bacterial burdens and M-MDSC infiltration in Mtb-immunized and BCG-infected mice. **A.** Schemes showing the genetic and pharmacological interference with M-MDSC. Mice remained unvaccinated or were immunized s.c. with 10^6^ CFU heat-killed Mtb/IFA at day 0 followed by a booster dose at day 12. On day 33, all mice were infected i.n. with 10^7^ CFU live BCG. Ccr2-DTR-CFP mice were treated with DT before or after live BCG infection. Similarly, C57BL/6 mice were treated with ATRA or placebo pellets before or after live BCG infection. Lungs and spleens were harvested and analyzed at day 47. **B.** Bacterial burdens in whole lungs and spleens. **C.** Frequencies of single cells in specified gates of lung and spleen. Gates as in Suppl. Fig. 1A. **B:** Statistics by unpaired, two-tailed t-test. n=4 biological replicates, 2 independent experiments. **C:** Statistics by ordinary one-way ANOVA with multiple comparisons and Tukey’s post test. n=3-4, 2 independent experiments. * p<0.05; ** p<0.01; *** p<0.001

To confirm the beneficial effect of genetic M-MDSC depletion, we next employed the pharmacological MDSC inhibitor ATRA ^37^ before and after BCG infection (Figure 5A). Similar to CCR2^+^ cell depletion, ATRA treatment led to a significant reduction in the bacterial loads in the lung and spleen (Figure 5B). Since the major effects of DT and ATRA on the bacterial loads were observed in the lung, we tested whether this correlated with the successful depletion of M-MDSC and how other myeloid cells were affected. As expected, M-MDSC declined after genetic or pharmacological interference with MDSC, along with monocytes, but not MoDC, Mph, or granulocytes (Figure 5C). Then, we investigated whether genetic and pharmacological interference with M-MDSC induction promotes cDC and T cell responses. The results indicate that cDC1 selectively increased at the early interference time points (Figure 6A). This correlated with elevated CD4^+^ T cell frequencies (Figure 6B) and higher expression of the activation markers CD69 and CD44 on the CD4^+^ T cell subset, while CD8^+^ T cells showed higher expression of the cytotoxicity-related marker Lamp1 (Figure 6C, D). Moreover, sera analysis of mice treated with ATRA showed elevated concentrations of inflammatory cytokines (Supplementary Figure 4). These data align with the results above, which show that MDSC depletion increased myeloid cell and T cell frequencies, activation, and function. Together, the depletion experiments support the concept that interfering with M-MDSC shortly after their induction by 2xMtb vaccination further lowers the bacterial loads by promoting cDC1, resulting in better expansion and activation of CD4^+^ and CD8^+^ T cells.

**Figure 6:**
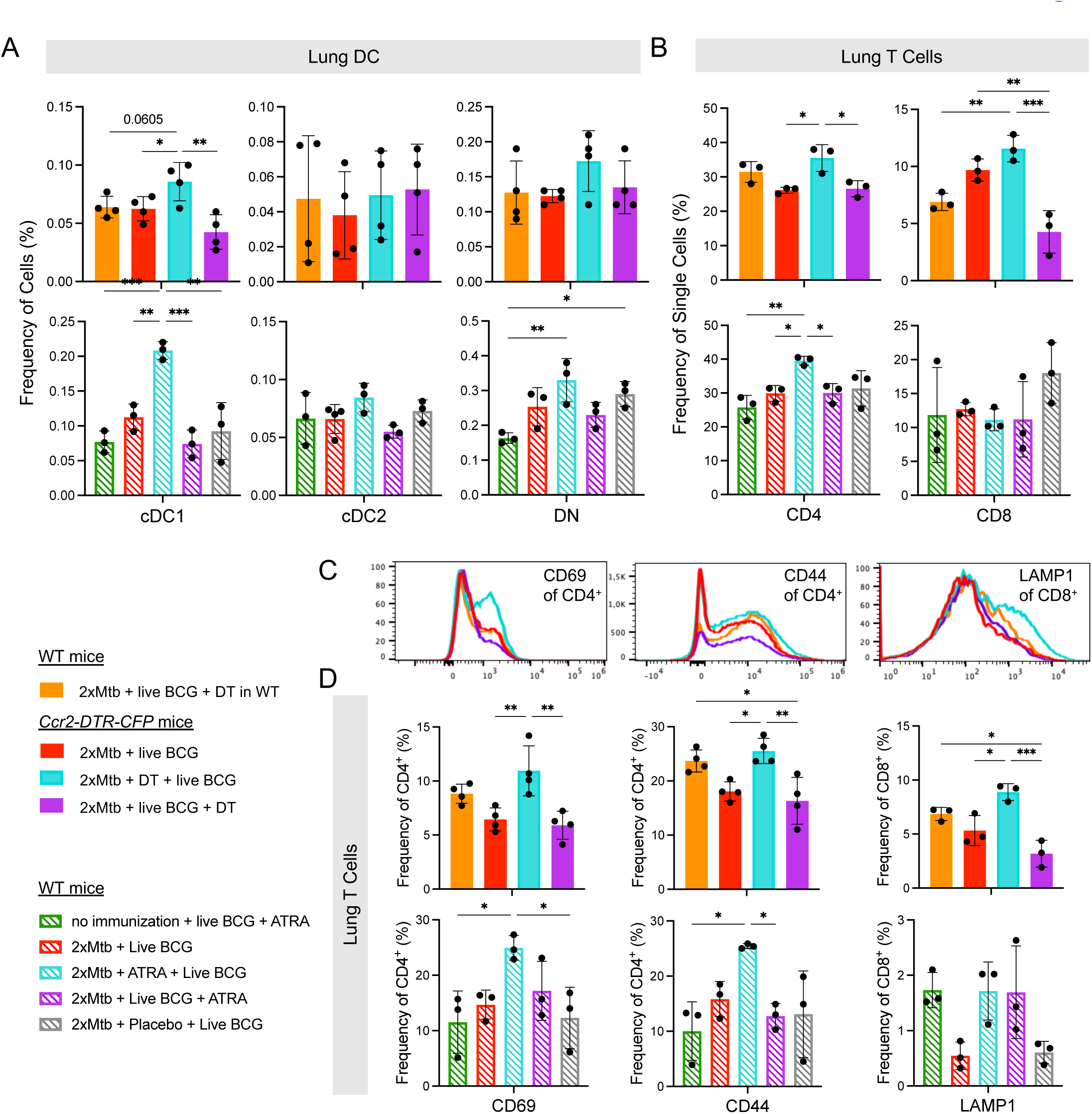
DT or ATRA treatment prior to intranasal BCG infection increases frequency of cDC1 and enhances activation of T cells in the lung. *Ccr2*-DTR-CFP or WT mice were immunized, infected and treated according the scheme shown in Fig. 5A. **A.** Frequencies of cells in specified DC populations in lung. Gates as in Fig. 4A. **B.** Frequencies of single cells in the CD4 and CD8 T cell gates. Gates as in Fig. 5A. **C.** Representative histograms showing expression of CD69 and CD44 activation markers in the CD4^+^ T cell gate and of LAMP1 in the CD8^+^ T cell gate from indicated conditions. **D.** Frequencies of CD69^+^ and CD44^+^ cells in CD4^+^ T cell gate and LAMP1^+^ cells in the CD8^+^ T cell gate. Statistics by ordinary one-way ANOVA with multiple comparisons and Tukey’s post test. n=3-4, 2 independent experiments. * p<0.05; ** p<0.01; *** p<0.001

## Discussion

Our previous data indicated that immunizing mice twice with heat-killed Mtb in IFA massively induced M-MDSC in the spleen, which entered the white pulp upon a microbial challenge to kill DC, thereby interfering with the induction of an immune response ^15^. Here, we used the same immunization (2xMtb) to extend our findings by vaccination with additional bacterial strains. These results indicated that qualitative differences exist between bacterial species in expanding M-MDSC. The question remains whether specific molecules expressed by some bacteria, such as Mtb, are responsible for M-MDSC expansion. There are major genetic differences between Mtb and BCG, most prominently known as ‘regions of difference’ (RD) ^38^. The RD1 region, present in virulent Mtb and Mbov but absent in avirulent BCG ^39^, has received the most attention, especially because it encodes the secreted molecules ESAT-6 and CFP-10, which represent major virulence factors and important antigens for T and B cells ^5^. However, no role for RD1 in MDSC induction has been shown, and we did not see differences between BCG and Mbov in their capacity to induce M-MDSC. The MTB64 protein of Mtb, also present in Mbov but not BCG, has been proposed to induce murine MDSC-like cells *in vitro* ^40^, which awaits confirmation *in vivo*. Thus, the question of whether a specific Mtb molecule can induce MDSC is still open. However, the fact that several different bacterial vaccines could induce MDSC here argues for a more general common mechanism.

Induction of MDSC may be driven by host factors such as inflammation or damage-associated molecular patterns to protect tissue integrity. The number of M-MDSC in secondary lymphoid organs or at infection sites will control the strength of immune responses that can occur locally. Increased frequencies of M-MDSC would allow a stronger control of immune responses. We showed previously that repetitive injections of GM-CSF into mice induced similar splenomegaly and M-MDSC induction as seen here after 2xMtb injection ^41^. It is tempting to speculate whether M-MDSC induction requires GM-CSF and whether its production is promoted better by 2xMtb injection than by the other bacteria tested here.

While 2xMtb immunizations led to massive M-MDSC induction in both lung and spleen. However, after BCG infection in these mice, we observed a decreased bacterial load in the lung but an increase in the spleen. The lower bacterial load in the lung correlated with higher frequencies and activation levels of several DC subsets and T cells than compared with the spleen. Thus, the local lung infection attracted both pro- and anti-inflammatory cells. The suppressor activity of M-MDSC isolated from these lungs indicated a parallel competing stimulation and inhibition of T cell responses. In our case, the dominant stimulatory response led to a lower bacterial load. Such competition between infiltrating pro- and anti-inflammatory cell types is well documented in tumors where infiltrating DC and T cells are controlled by MDSC, frequently tipping the balance towards more tumor growth ^9^ ^42^. Similarly here, high frequencies of M-MDSC were already present in the spleen when BCG disseminated to this organ, and thereby suppression was dominating.

Surprisingly, the 2xBCG heat-killed vaccination showed neither signs of splenomegaly nor an innate or adaptive immune response. Lung and spleen showed even a trend towards higher bacterial loads than the controls. We used the same dose of heat-killed BCG to compare with the established 2xMtb immunization scheme for M-MDSC induction. It has been previously observed in humans that there was no significant change in control of BCG outgrowth after revaccination with BCG ^43^. Alternatively, the live BCG vaccine has been shown to be more immunogenic than the killed bacteria ^44,45^, which may be due to a longer persistence, possibility of dissemination and increased tissue damage, rarely observed during states of immunodeficiency ^46,47^.

While TB vaccination with BCG is widely used, approval for a Mtb-based vaccine is still pending. Mainly, superior immune evasion mechanisms of Mtb have been discussed for this failure ^48,49^. To improve the vaccination success against TB, adjunct host-directed therapies have been proposed ^21,50^. Here, we tested whether the successful Mtb vaccination protecting from BCG infection could be further improved when interfering with M-MDSC. While previous studies demonstrated that ATRA can treat Mtb-infected mice successfully ^8,23,24^, we show here that it can also be used as an adjunctive treatment with an Mtb-based vaccine. At early time points (d15/16/17) when the maximal M-MDSC induction could be observed, both CCR2^+^ cell-depleting genetic approach and the pharmacological treatment with ATRA diminished frequencies and function of M-MDSC, the latter as reported after treatment of tumor-bearing mice ^22^, together with elevated inflammatory cytokines and higher frequencies of cDC1 and CD4^+^ T cells in the lung. Interference at later time points (d34/35/36), after live BCG infection, failed to reduce bacterial counts, enhance DC numbers, or improve T cell responses despite reducing the M-MDSC frequencies in the lung. While DT depletion of CCR2^+^ cells and their descendants affects pro-inflammatory MoDC and suppressive M-MDSC, ATRA is believed to affect only MDSC. Since the genetic and pharmacological approaches behaved very similarly here, this does, however, suggest that ATRA does not exclusively act on suppressive cells but also inhibits the differentiation of pro-inflammatory cells such as DC. This seems to contrast with what has been observed in human studies where DC vaccination against small cell lung cancer was successfully combined with ATRA treatment ^28^. However, the vaccines in this study were already mature DC, which did not require differentiation from progenitors or monocytes. ERK1/2 signaling, a major pathway responsible for ATRA-mediated inhibition of MDSC differentiation ^22^, plays an important role in DC survival but not maturation ^51^. Thus, the optimal time point to interfere with M-MDSC in a vaccine setting has to occur in conjunction with the vaccine but not after infection, where pro-inflammatory DCs are generated from progenitors or monocytes. Recent clinical studies to interfere with MDSC by ATRA as an adjunct treatment with checkpoint inhibitors of tumor patients revealed encouraging results^27,52^.

In conclusion, our work provides several new insights into mycobacterial vaccination. First, 2xMtb immunization showed the highest potential to induce M-MDSC over the other bacteria tested. Second, despite the induction of M-MDSC after 2xMtb vaccination, it also induced a substantial immune response that lowered the bacterial loads in our setting. Third, ATRA treatment during the 2xMtb vaccination to interfere with M-MDSC further improved the immune response and protection from infection. Together, these results may encourage the future use of ATRA in human vaccination studies against TB and other pathogens where MDSC induction has been observed ^53^.

## Materials and Methods

### Mice and interference with M-MDSC

P25 transgenic mice ^36^ were kindly provided by Ulrich Schaible, Borstel, Germany. P25 mice were crossed with B6.CD45.1 congenic mice to obtain B6.P25.CD45.1 mice. C57BL/6J wild-type mice were initially purchased from Charles River, Sulzfeld, Germany. All mice were bred in our animal facilities and kept under specific pathogen-free conditions. Both sexes were used for experiments at an age of 6-12 weeks. Georg Gasteiger kindly provided Ccr2-DTR-CFP mice. Diphtheria toxin (DT, Unnicked, *Corynebacterium diphtheriae*, Merck, Darmstadt, Germany) was injected i.p. at indicated time points at a dose of 0.5µg per injection in PBS. Pellets containing all-trans retinoic acid (ATRA) or equivalent placebo were applied s.c. (ATRA 5mg/p, 21-day release, Innovative Research of America, Sarasota, FL) at the indicated time points. Before BCG infection and ATRA implantation, all mice were anaesthesized by i.p. injection of Ketamine (100µg/g) and Medetomidine (1µg/g) mixture in 0.9% NaCl solution. Finally, as a reversal of anesthesia, mice were inoculated i.p. with 12.5µg/g Anti-sedan (active ingredient Atipamezole Hydrochloride) in 0.9% NaCl solution. The euthanization of animals was performed in a CO_2_ chamber. All animal euthanizations, treatments, and experiments were performed according to German animal protection law and after approval and under the control of the local authorities (Regierung von Unterfranken, AZ 55.2-2531.01-64/11 and 55.2.2-2532-2-1408).

### Bacterial immunization

Heat-killed bacteria were mixed in Incomplete Freund’s Adjuvant (IFA, purchased from MilliporeSigma) or aluminium hydroxide (alum, purchased from Brenntag) and emulsified in an equal amount of PBS. A creamy preparation was achieved by passing the mixture continuously through 2 syringes connected by a medical valve 3-way stopcock for 10 minutes. C57BL/6J mice were immunized s.c. by injecting 200µl bacteria/IFA-PBS or, where indicated, bacteria/alum-PBS emulsion into one flank. Booster immunization was administered s.c. by introducing 200μl emulsion into the other flank. When indicated, 10^7^ bulk spleen cells from B6.P25.CD45.1 mice in a total volume of 50µl per mouse were injected i.v. into mice tail veins.

### Bacteria and infection

Heat-killed and lyophilized Mtb H37Ra was purchased from Difco (Mtb). BCG Pasteur expressing red fluorescence (dsRed) (BCG) was kindly provided by Nathalie Winter, Pasteur Institute Paris, France ^54^. *Mycobacterium smegmatis* (Msm) was kindly provided by Sebastian Geibel, Würzburg, Germany. Heat-killed *Listeria monocytogenes* (List) was provided by Thomas Hünig, Würzburg, Germany. Heat-killed but non-lyophilized H37Ra Mtb was prepared by our Microbiology Department (MtbM). MtbM, BCG, Msm, and wild-type Mbov were cultured, heat-inactivated, and prepared for immunization in-house, all in the same way. Both Mtb preparations used in this study were of the H37Ra strain. All mycobacteria were cultured in Middlebrook 7H9 broth medium supplemented with 0.05% Tween-80, 0.05% glycerol, 10% ADC (Albumin, Dextrose, Catalase), and 30µg/ml Hygromycin at 35°C until reaching exponential growth phase. The bacterial suspension was then centrifuged at 30 G for 10 minutes, quantified, used, or stored in 20% glycerol at −20°C. When necessary, BCG was heat-killed by placing aliquots in a water bath at 80°C for 60 minutes. Infection was performed by i.n. bacterial inoculation of 20µl droplets (5 x 10^8^CFU/ml).

### Evaluation of infection by determination of bacterial load

Lung and spleen bacterial loads of BCG-infected mice were determined by plating serial dilutions of whole-organ homogenates on Middlebrook 7H11 agar medium enriched with ADC. Petri dishes were covered with permeable parafilm and incubated at 37°C in humidified air containing 5% CO_2_. Bacterial colonies were counted 2-3 weeks post-incubation.

### Flow cytometry

The murine directly conjugated antibodies CD11b-Alexa Fluor 700 / -Brilliant Violet 650 (M1/70), Ly6G-Brilliant Violet 650 / -APC-Cy7 (1A8), Ly6C-Brilliant Violet 510 / -Brilliant Violet 785 (HK1.4), CD11c-PE-Cy7 (N418), CD16-2-Brilliant Violet 421 (9E9), B220-Pacific Blue (RA3-6B2), XCR1-APC (ZET), CD64-PE (X54-5/7.1), MHC II-Alexa Fluor 700 (M5/114.15.2), CD4-PerCPCy5.5 / -Pacific Blue (GK1.5), CD8-Alexa Fluor 488 / -Brilliant Violet 510 (53-6.7), Vβ11-APC (RR3-15), CD45.1-Pacific Blue (A20), Ki67-PE (11F6), CD69-PE-Cy7 (H1.2F3), CD44-Alexa Fluor 488 (IM7), LAMP1/CD107a-PE (1D4B), PD-L1/CD274-PE (10F.9G2), Annexin V-FITC were all purchased from Biolegend. S100A9-APC (2B10) was bought from BD Pharmingen. iNOS-FITC (CXNFT), Arg1-APC (A1exF5), Fixable Viability Dye-eFluor 780 (#65-0865) were obtained from eBioscience. Surface staining was performed by incubating 10^6^ cells in 100µl FACS buffer containing fluorescent-conjugated antibodies for 30 minutes at 4°C. To block unspecific antibody binding to surface FcγRII/III, 50% supernatant from the 2.4G2 hybridoma was used in FACS buffer. Cells were then fixed with 2% formaldehyde for 20 minutes at room temperature. For intracellular iNOS and Arg1 detection, cells were permeabilized and stained in 100µl 1x Perm buffer with fluorescent-labeled antibodies for 1 hour at room temperature. Samples were washed with Perm buffer, resuspended in FACS buffer, and measured with the Attune NxT (Thermo Fisher Scientific). Data were analyzed by FlowJo 10 (Tree Star).

### Activation of M-MDSC

M-MDSC were activated *ex vivo* to induce iNOS or Arg1 expression. Spleens and lung cell suspensions were seeded in 24-well plates at a density of 1 x 10^6^ cells/ml per well. They were cultured in the presence of 100ng/ml Lipopolysaccharide (LPS, MilliporeSigma) and 100U/ml IFN-γ (Immunotools) for 16 hours as described ^12,41^.

### Suppressor Assay

Bulk spleen and lung single-cell suspensions were seeded in 96-well plates at a concentration of 2 x 10^5^ cells/well in 200µl RPMI medium containing 10% FCS and stimulated with soluble anti-CD3/anti-CD28 antibodies at a final concentration of 2.5µg/ml each. In P25 spleen cell transfer experiments, cells were stimulated with 10µg/ml P25 peptide (FQDAYNAAGGHNAVF) instead. At day 3, proliferation was measured by flow cytometry via Ki67 detection separately in CD4^+^ and CD8^+^ T cell subsets. iNOS inhibitor N^G^-Methyl-L-arginine acetate salt (L-NMMA, 500 μM, MilliporeSigma) and Arg1 inhibitor Nω-hydroxy-nor-arginine (nor-NOHA, 50μg/ml, Cayman Chemicals) were added to the cultures when required.

### Legendplex Analysis

On the day of sacrifice, blood was collected by cardiac puncture, and the serum was separated by centrifugation at 3.6G for 30 minutes at 4°C. The samples were stored at -20°C until thawed and tested for the presence of cytokines using the LEGENDplex™ Mouse Inflammation Panel 13-plex kit (Biolegend) following the manufacturer’s protocol. The samples were then measured with the Attune NxT (Thermo Fisher Scientific) and analyzed using the LEGENDplex™ online software (Biolegend).

### Statistics

Figures were created, and statistics were calculated using GraphPad Prism 10 software. Details on the statistical tests used are provided in the figure legends and include student’s t-test, one-way or two-way ANOVA as indicated. P values of less than 0.05 were considered significant.

## Acknowledgments

We thank Sabrina Schneider and Marion Heuer for their expert technical assistance. This work was supported by funding for AA and MBL through the German Research Foundation (DFG LU851/18-1). HA is supported by a scholarship from the Ministry of Higher Education and Scientific Research of the Arabic Republic of Egypt.

## Author contributions

AA, HA, LC performed the experiments, analyzed the data and prepared the figures; CS, US provided valuable reagents and mice; AA, NDP, GW and MBL conceptualized the work and together with NN interpreted the data; AA, NN and MBL wrote the paper.

## Abbreviations

M-MDSC: Monocytic myeloid-derived suppressor cells
TB: Tuberculosis
Mtb: *Mycobacterium tuberculosis*
BCG: Bacille-Calmette-Guérin
Mbov: *Mycobacterium bovis*
Msm: *Mycobacterium smegmatis*
List: *Listeria monocytogenes*
IFA: Incomplete Freund’s Adjuvant
CFA: Complete Freund’s Adjuvant
LPS: Lipopolysaccharide
DT: Diphtheria Toxin
ATRA: All-trans retinoic acid

