## Supplemental data for "MDSC depletion during immunization with heat-killed *Mycobacterium tuberculosis* increases protection against BCG infection"

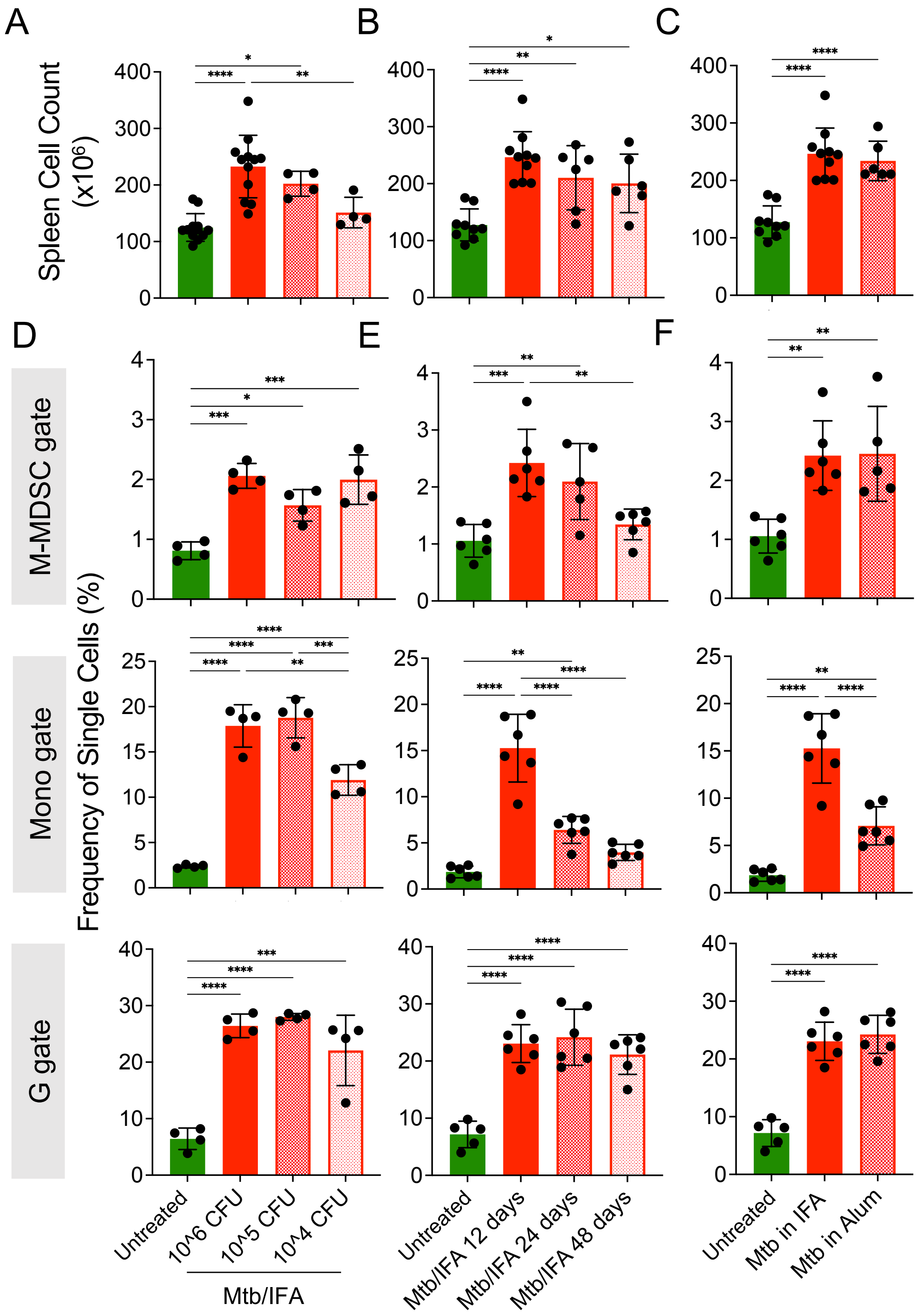


**Suppl. Fig. 1: Minor differences in M-MDSC induction after Mtb immunization with variations in bacterial dose, time interval or type of adjuvant.**

**A-F.** The standard protocol was compared to immunizations with titrated Mtb doses (**A, D**), extended prime-boost immunization intervals (**B, E**), or Alum adjuvant (**C, F**). Spleen cell numbers **(A.B,C)** or the frequency of M-MDSC, monocytes and granulocytes among single cells **(D, E, F)** were measured. Statistics by ordinary one-way ANOVA with multiple comparisons and Tukey's post test. n=4-12 biological replicates, 2-6 independent experiments. * $p<0.05$; ** $p<0.01$; *** $p<0.001$; **** $p<0.0001$

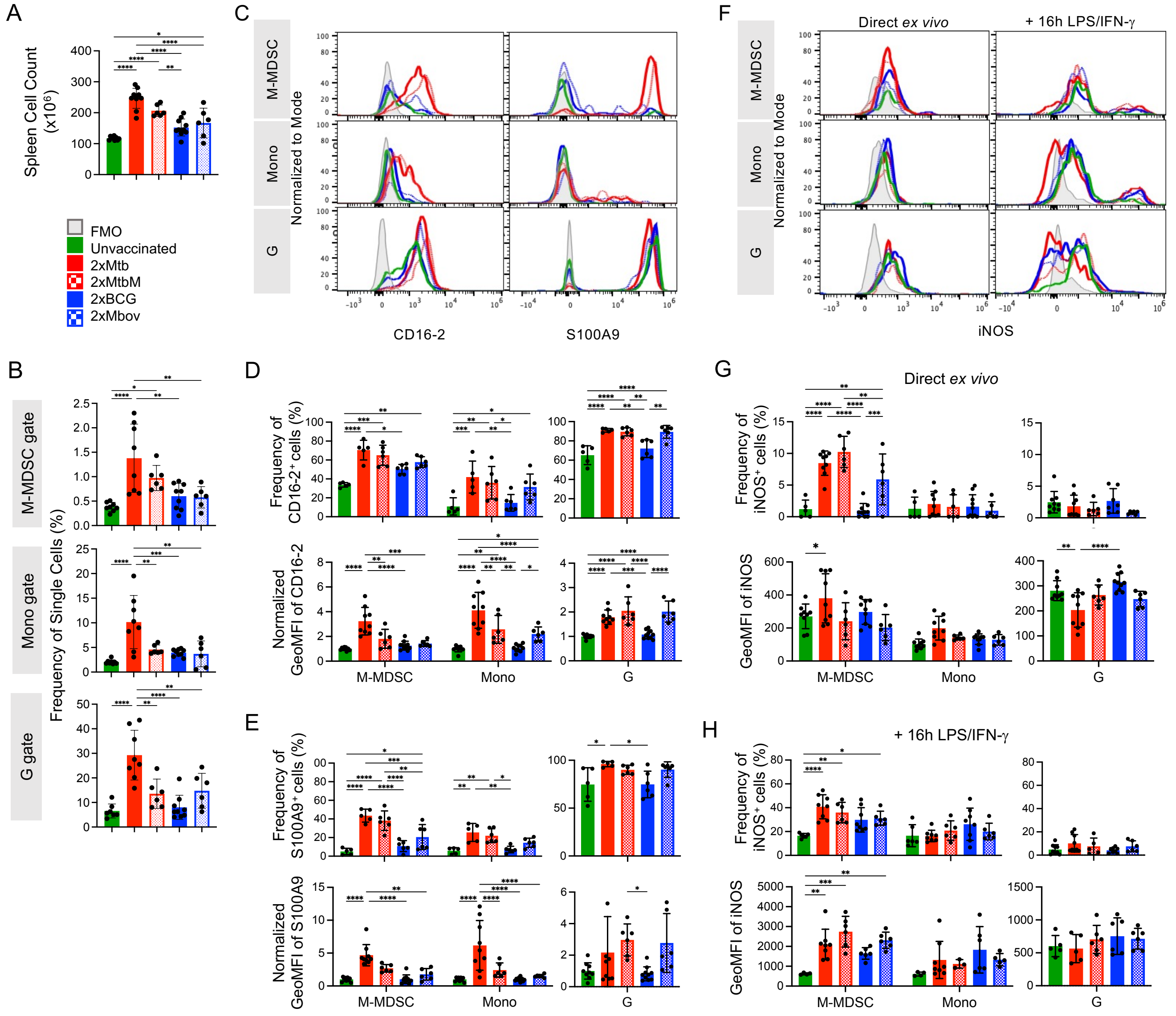

**Suppl. Fig. 2: Comparison of immunization with lyophilized versus non-lyophilized Mtb and BCG versus Mbov.**

Mice remained unvaccinated or were immunized s.c. with heat-killed and lyophilized Mtb (Mtb) in IFA, heat-killed but non-yophilized Mtb (MtbM) in IFA, heat-killed BCG in IFA or heat-killed Mbov in IFA as indicated at day 0 followed by a booster dose at day 12, and spleens were analyzed at day 15. **A.** Spleen cell numbers were determined. Unvaccinated and 2xMtb groups same as in Fig. 1B for comparison. **B.** Frequencies of single cells in M-MDSC, monocytes (Mono) and granulocytes (G). Gates as in Fig. 1C. **C.** Representative histograms showing expression of CD16-2 and S100A9 activation markers in M-MDSC, Mono and G gates. **D, E.** Frequency and normalized GeoMFI of CD16-2 and S100A9 in M-MDSC, Mono and G gates, respectively**.** **F.** Spleen cells were restimulated *in vitro* by LPS/IFN-γ for 16h then stained for iNOS. Histograms showing iNOS expression directly *ex vivo* and after restimulation *in vitro* in M-MDSC, Mono and G gates. **G, H.** Frequency and GeoMFI of iNOS in M-MDSC, Mono and G gates directly *ex vivo* and after restimulation, respectively. **A, B:** Statistics by ordinary one-way ANOVA with multiple comparisons and Tukey's post test. n=6-10 biological replicates, 2-4 independent experiments. **D, E, G, H:** Statistics by two-way ANOVA with multiple comparisons and Tukey's post test. n=6-10 biological replicates, 2-4 independent experiments. * p<0.05; ** p<0.01; *** p<0.001; **** p<0.0001

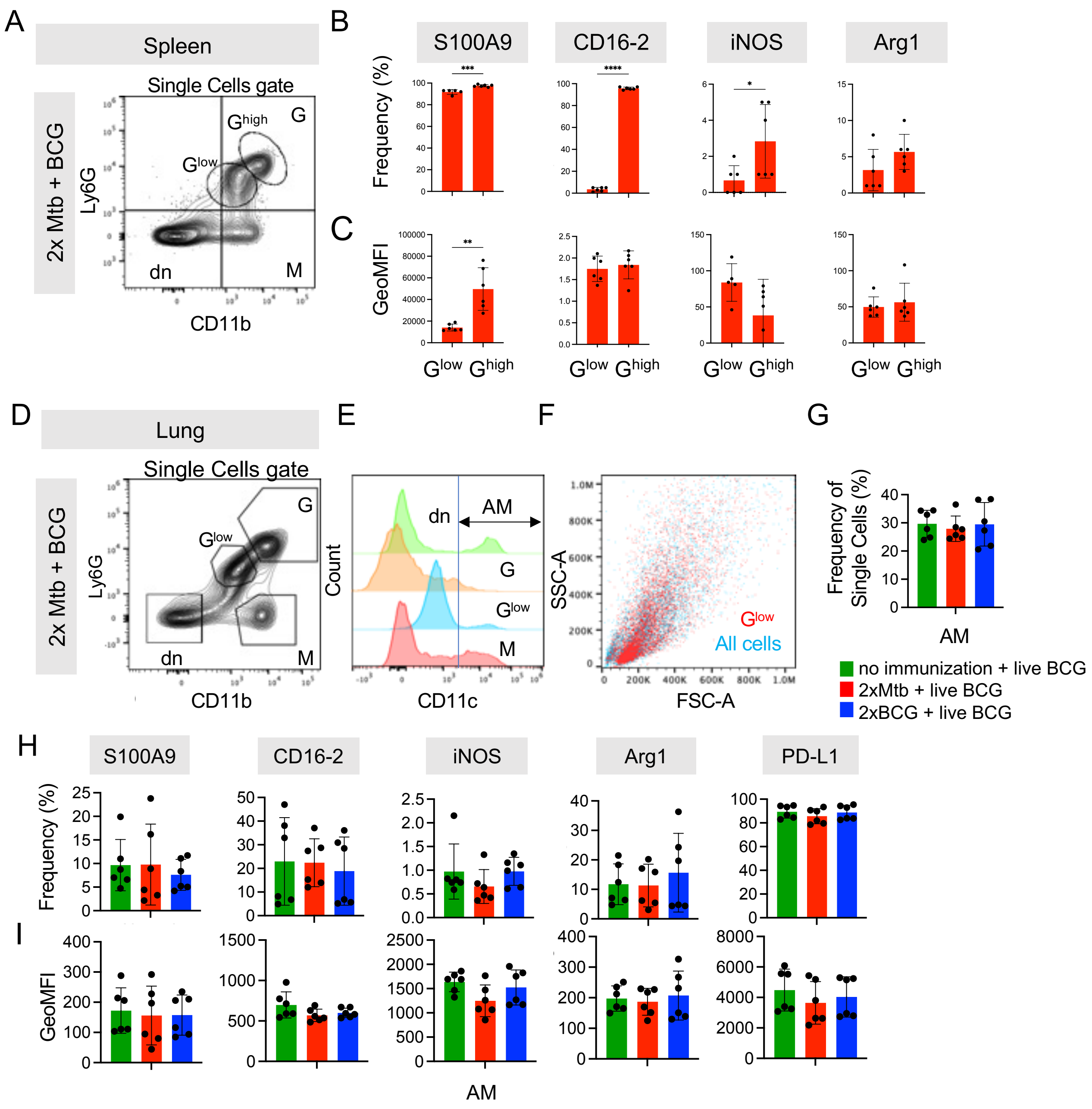


**Suppl. Fig. 3: Lung granulocyte subset analysis points towards two neutrophil maturation stages and alveolar macrophages remain functionally unaffected by immunizations.**

**A.** Flow cytometry example plot of single cell gate from mouse spleen after infection. Spleen G gate was further divided into $G^{low}$ ($Ly6G^{low}$ $CD11b^{low}$) and $G^{high}$ ($Ly6G^{high}$ $CD11b^{+}$). **B, C.** Frequencies and GeoMFI of indicated activation and functional markers in spleen $G^{low}$ and $G^{high}$ gates of 2xMtb + BCG infected mice. Gates as in A. **D.** Representative plot showing the gating strategy for the mouse lung in the flow cytometry analysis. **E.** CD11c expression of cells as gated in D of 2xMtb + BCG infected mouse. Arbitrary blue line separates low versus high CD11c expressing cells. Alveolar macrophages (AM) are designated as $CD11c^{+}$ within Ly6G and CD11b double negative (dn) gate. **F.** Flow cytometry plot showing the overlay of the $G^{low}$ gate (in red) on all cells (in blue). **G.** Frequency of single cells in the AM gate. Gate as in E. **H, I.** Frequencies and GeoMFI of indicated activation and functional markers in lung AM gate, under different immunization conditions followed by BCG infection. **B, C:** Statistics by unpaired, two-tailed t test. n=12 biological replicates pooled 2 by 2 (therefore n=6), 2 independent experiments. **G-I:** Statistics by ordinary one-way ANOVA with multiple comparisons and Tukey's post test. n=12 biological replicates pooled 2 by 2 in 2 independent experiments. * $p<0.05$; ** $p<0.01$; *** $p<0.001$; **** $p<0.0001$

### Suppl. Fig. 4

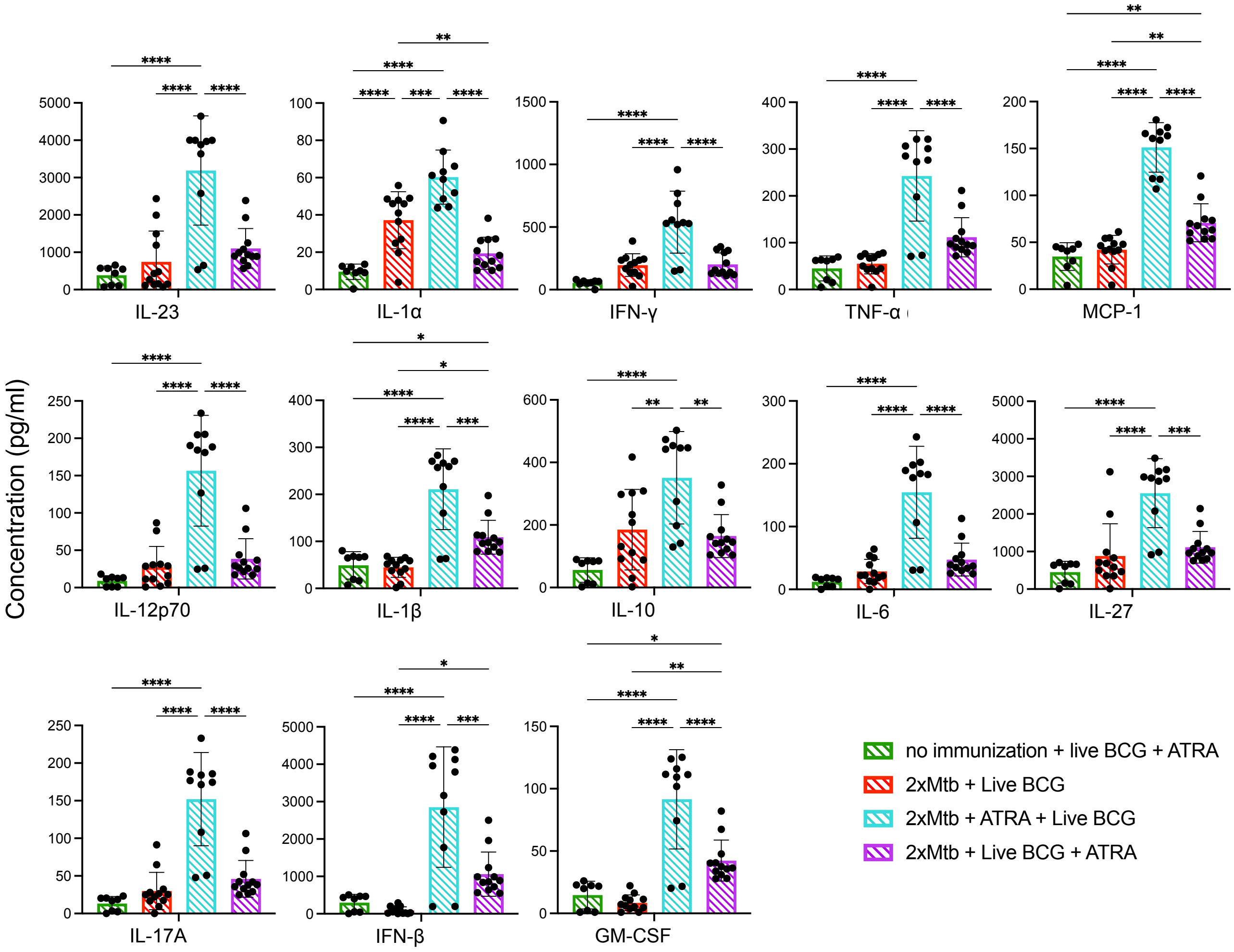


**Suppl. Fig. 4: ATRA treatment before intranasal BCG infection increases the concentration of inflammatory cytokines in mouse blood.** WT mice were immunized and infected according to the scheme shown in Fig. 5A. Blood was collected on day 47, and serum was analyzed by Legendplex analysis for the indicated cytokines. Statistics by ordinary one-way ANOVA with multiple comparisons and Tukey's post-test. n=12 biological replicates pooled 2 by 2 in 2 independent experiments. * $p<0.05$; ** $p<0.01$; *** $p<0.001$; **** $p<0.0001$
